# Targeting ATR-CHK1-WEE1 signaling enhances IMPDH inhibitor efficacy in mTORC1-driven tumor cells

**DOI:** 10.64898/2026.11.23.768592

**Authors:** Masuda Akther, Alexandra Grumet, Rayees A. Padder, Jay N. Joshi, Nanda Singh, Joonyoung Her, Samuel F. Bunting, Alexander J. Valvezan

## Abstract

Uncontrolled activation of the master metabolic regulator Mechanistic Target of Rapamycin Complex 1 (mTORC1) in tumors confers strong dependence on the guanylate nucleotide synthesis enzyme inosine-5′-monophosphate dehydrogenase (IMPDH) for tumor cell growth and viability. Clinically approved IMPDH inhibitors reduce the guanylate nucleotide pool available for DNA synthesis in mTORC1-driven tumor cells, resulting in prolonged DNA replication stress and accumulation of DNA damage that culminates in apoptosis. These early molecular events are accompanied by activation of adaptive DNA replication stress response (RSR) and DNA damage response (DDR) pathways. We hypothesized that targeting these adaptive responses could enhance IMPDH inhibitor efficacy. We find that combining IMPDH inhibition with inhibition of the ATR-CHK1-WEE1 signaling axis increases the efficacy and durability of response in cell models, and in a syngeneic xenograft tumor model of the mTORC1-driven genetic tumor syndrome Tuberous Sclerosis Complex. Mechanistically, abrogating ATR-CHK1-WEE1 signaling during prolonged IMPDH inhibition causes aberrant cell cycle progression, increased DNA double-strand breaks, and formation of micronuclei. Together these findings uncover adaptive mechanisms that restrict IMPDH inhibitor efficacy, informing a combination strategy to improve treatment efficacy, while maintaining selectivity for targeting tumor cells that are driven by uncontrolled mTORC1 activation.

## Introduction

Proliferating cells must balance biosynthesis with nutrient availability for sustainable growth. Mechanistic Target of Rapamycin Complex 1 (mTORC1) is a master metabolic regulator that integrates nutrient and growth factor signals to control the balance between anabolic and catabolic processes. Under growth-promoting conditions, mTORC1 coordinates an extensive anabolic network to synthesize the major macromolecules required for cell growth, including proteins, lipids, and nucleic acids, while inhibiting catabolic processes such as autophagy (1, 2). mTORC1 also stimulates flux through metabolic pathways that support macromolecule synthesis, including the *de novo* purine and pyrimidine nucleotide synthesis pathways, glycolysis, and the oxidative branch of the pentose phosphate pathway (3–6). Owing to its central importance in promoting anabolic growth, mTORC1 is activated in the majority of human cancers across nearly all lineages (7, 8). This occurs downstream of many of the most commonly mutated oncogenes and tumor suppressors in human cancer, including receptor tyrosine kinases, *PI3KCA, PTEN, AKT, RAS, RAF, APC*, and others (7, 9). Oncogenic mutations in these upstream regulators converge to inhibit the TSC complex, an essential negative regulator of mTORC1, resulting in uncontrolled mTORC1 activation (7, 9).

Loss-of-function mutations in the TSC complex components *TSC1* and *TSC2* cause the genetic tumor syndrome Tuberous Sclerosis Complex (TSC), in which tumor growth is driven by mTORC1 hyperactivation (10, 11). The mTORC1 inhibitor rapamycin, and its analogs, are the current standard of care therapeutics for treating TSC tumors. However, these agents are partial, allosteric inhibitors of mTORC1 that generally induce cytostatic responses by downregulating anabolic pathways, rather than cytotoxic responses, in TSC tumors and in cancer (12–14). Thus, tumors are not eliminated and resume growth when treatment is discontinued. Consequently, TSC patients must undergo lifelong treatment, underscoring the need for improved therapeutic approaches to selectively kill mTORC1-driven tumor cells.

As an alternative approach to targeting mTORC1 itself, recent understanding of the coordinated anabolic network activated downstream of mTORC1 has opened the door for targeting metabolic vulnerabilities, i.e. metabolic pathways that become uniquely essential in the context of uncontrolled mTORC1 activation. Exploiting such vulnerabilities is advantageous in that it can imbalance the mTORC1-driven anabolic program, causing accumulation of cellular stresses, which culminate in selective tumor cell death *in vitro* and *in vivo* (15–19). We found that uncontrolled mTORC1 activation in cell and tumor models of TSC, or downstream of PTEN-loss in T-cell acute lymphoblastic leukemia, greatly increases the cellular demand for nucleotides by driving the synthesis of ribosomal RNA (rRNA) for ribosome biogenesis. This creates strong dependence on *de novo* nucleotide synthesis pathways, which are stimulated by mTORC1 in parallel, to sustain sufficient free NTP pools. Targeting *de novo* nucleotide synthesis pathways causes selective nucleotide depletion in tumor cells that have uncontrolled mTORC1 activation because of their highly elevated nucleotide demand, resulting in prolonged and selective DNA replication stress, accumulation of DNA damage, and ultimately apoptotic cell death (15–17).

This unique metabolic dependency can be exploited using clinically approved inhibitors of the rate-limiting enzyme in *de novo* guanylate nucleotide synthesis, inosine 5’-monophosphate dehydrogenase (IMPDH) (15–17). IMPDH inhibitors have been used for decades as safe and well-tolerated immunosuppressants (20, 21) but are not currently approved as anti-tumor agents. IMPDH has also emerged as a promising target for anti-cancer therapy in preclinical models of small-cell lung cancer (22, 23), hepatocellular carcinoma (24–26), leukemia (17, 27, 28), and glioblastoma (29–31), with 2 clinical trials in glioblastoma currently in progress (NCT05236036 and NCT04477200).

Direct comparison of the IMPDH inhibitor mizoribine (Bredinin) with rapamycin in TSC tumor models demonstrates that mizoribine can produce more durable tumor control due to its ability to induce DNA replication stress, DNA damage, and tumor cell death, in contrast to rapamycin (15, 16). However, despite strong initial efficacy, tumors treated with mizoribine are not completely eliminated and they eventually regrow (albeit slowly) when treatment is discontinued. Thus, we sought to further enhance the efficacy and durability of IMPDH inhibitor responses.

Given the mechanism of action of IMPDH inhibitors, early cellular responses are accompanied by activation of adaptive DNA replication stress response (RSR) and DNA damage response (DDR) pathways (15–17). We hypothesized that targeting these adaptive responses could enhance IMPDH inhibitor efficacy. RSR/DDR pathways are mediated by serine/threonine protein kinases in the PI3K-Related Kinase (PIKK) family, including ATR, ATM, and DNA-PK, to activate cell cycle checkpoints, allow recovery from replication stress, and repair DNA damage (32, 33). Many of these functions are mediated by the CHK1 and CHK2 effector proteins, which are phosphorylated and activated by ATR and ATM, respectively. Activated CHK1 then activates WEE1 in a linear pathway that arrests cells at the G2/M checkpoint to prevent cells from entering mitosis with damaged or under-replicated DNA (34, 35). Together, these mechanisms promote cell survival under conditions of DNA replication stress and DNA damage. Numerous clinical trials are currently being conducted with inhibitors targeting these RSR/DDR pathways in cancer (36–39).

We investigated whether pharmacological targeting of key RSR/DDR kinases would enhance the efficacy of IMPDH inhibitors, while maintaining selectivity by preferentially affecting cells that accumulate DNA replication stress and DNA damage upon IMPDH inhibition (i.e. the mTORC1-driven tumor cells). Using TSC2-deficient cells and tumors as models of mTORC1-driven growth, we demonstrate that inhibiting the ATR-CHK1-WEE1 signaling axis enhances IMPDH inhibitor-induced cell death and significantly slows regrowth when treatment is discontinued. Mechanistically, addition of ATR-CHK1-WEE1 inhibitors during prolonged IMPDH inhibition causes aberrant cell cycle progression, increased DNA double-strand breaks, and formation of micronuclei. Together these findings support combined inhibition of IMPDH and RSR/DDR kinases as a potential therapeutic strategy for effective tumor cell killing, with more durable treatment responses compared to single agent therapy or mTORC1 inhibition. Our use of inhibitors that are clinically approved or in advanced clinical development could facilitate translation of this approach to patients with TSC and cancers driven by uncontrolled mTORC1 activation.

## Results

### IMPDH inhibitor efficacy is enhanced in TSC2-deficient tumor cells by inhibition of ATR-CHK1-WEE1 signaling

IMPDH inhibition selectively induces DNA replication stress and DNA damage in cells with uncontrolled mTORC1 activation, ultimately leading to apoptosis (15–17). These early molecular events are accompanied by activation of replication stress response (RSR) and DNA damage response (DDR) pathways. We hypothesized that targeting these adaptive mechanisms could enhance IMPDH inhibitor efficacy. To test this, *Tsc2^×/×^*105K renal tumor cells were treated with the clinically approved IMPDH inhibitor mizoribine, alone or in combination with the ATR inhibitor AZD6738/ceralasertib. Colony formation was measured at the end of the treatment period, and in parallel plates after a 3-day regrowth period following drug washout. Combining mizoribine with AZD6738 reduced colony formation and prevented regrowth compared to either treatment alone (Figure 1A,B). Time course analysis of cell growth and viability confirmed that combined treatment with mizoribine plus AZD6738 reduced the number of viable cells and prevented their regrowth after drug washout, whereas cells treated with mizoribine alone resumed gradual proliferation (Figure 1C). To investigate the fate of cells that remain after initial treatment, senescence-associated β-galactosidase (SA-β-gal) activity was measured at the end of the regrowth period. Combination treatment increased the proportion of SA-β-gal positive cells, indicating that inhibition of IMPDH plus ATR induces senescence in surviving cells (Figure 1D,E).

**Figure 1.**
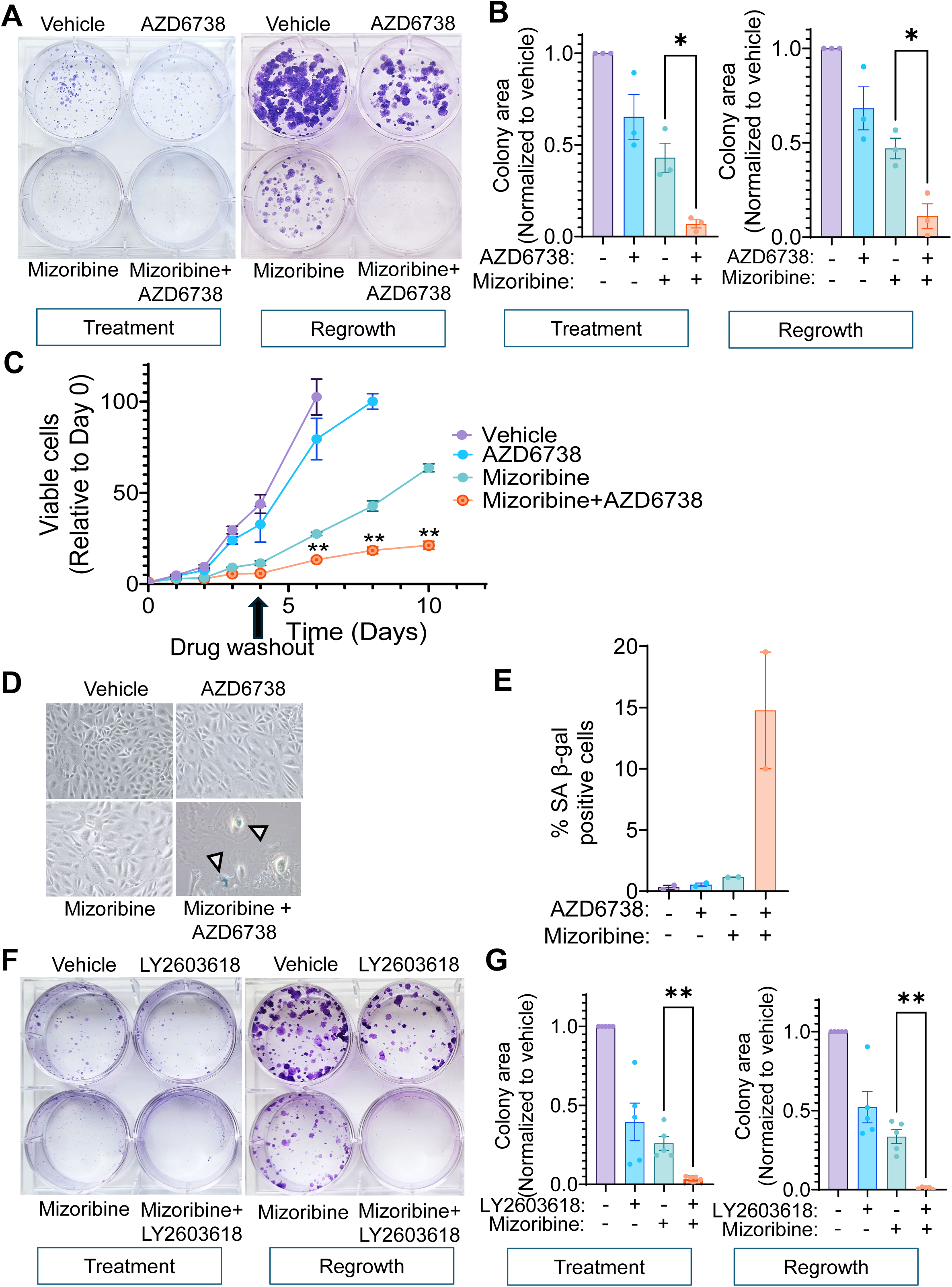
IMPDH inhibitor efficacy is enhanced by combined inhibition of ATR or CHK1. **(A, B)** *Tsc2^−/−^* 105K renal tumor cells were treated for 6 days with the IMPDH inhibitor mizoribine (5 µM) alone and in combination with the ATR inhibitor AZD6738 (1 µM). Following treatment, parallel plates were washed, replenished with fresh media, and allowed to regrow in the absence of drug treatments for an additional 3 days. **(A)** Colonies were visualized by crystal violet staining. **(B)** Colony area in (A) was quantified and graphed as fold change relative to the vehicle. **(C)** Cells in (A) were treated for 4 days with mizoribine (10 µM) alone and in combination with the ATR inhibitor AZD6738 (1 µM). On Day 4, cells were washed, replenished with fresh media, and allowed to regrow in the absence of drug treatments for up to 6 days. Viable cells were quantified using the CellTiter-Glo cell viability assay (Promega), and graphed normalized to day 0. **(D, E)** Cells in (A) were treated and allowed to regrow as described in (A, B). **(D)** Senescence-associated β-galactosidase (SA-β-gal) staining was performed at the end of the 3-day regrowth period. **(E)** SA-β-gal positive cells in (D) were quantified and graphed as percent of total cells. **(F, G)** Cells in (A) were treated for 6 days with mizoribine (3 µM) alone and in combination with the CHK1 inhibitor LY2603618 (0.25 µM). Following treatment, parallel plates were washed, replenished with fresh media, and allowed to regrow in the absence of drug treatments for an additional 3 days. **(F)** Colonies were visualized by crystal violet staining. **(G)** Colony area in (F) was quantified and graphed as fold change relative to the vehicle. Graphical data are presented as mean of at least three biological replicates ± SEM. *p<0.05, **p<0.006 by Welch’s t-test in (B,G), and by two-way ANOVA with Bonferroni’s multiple comparison test comparing mizoribine to mizoribine plus AZD6738 in (C).

To determine whether targeting other RSR/DDR kinases can similarly enhance IMPDH inhibitor efficacy, 105K cells were treated with mizoribine in combination with small-molecule inhibitors targeting CHK1 (LY2603618/rabusertib), CHK1/CHK2 (AZD7762), WEE1 (AZD1775/adavosertib), ATM (AZD0156), or DNA-PK (AZD7648). Co-inhibition of IMPDH and CHK1 (Figure 1F,G), CHK1/CHK2 (Supplemental Figure 1A,B), or WEE1 (Supplemental Figure 1C,D) impaired colony formation and/or prevented regrowth after drug washout compared to each inhibitor alone. In contrast, inhibition of ATM trended towards enhancing IMPDH inhibitor efficacy but did not reach statistical significance, and DNA-PK inhibition did not enhance efficacy (Supplemental Figure 1E–H). These data indicate that IMPDH inhibitor efficacy can be enhanced by combined inhibition of the linear ATR-CHK1-WEE1 signaling axis.

### RSR/DDR kinase inhibition enhances IMPDH inhibitor-induced apoptosis while maintaining selectivity for TSC2-deficient cells

IMPDH inhibition induces apoptotic cell death in TSC2-deficient cells (15, 16). To determine the effect of combined inhibition of IMPDH and RSR/DDR kinases, 105K cells, *Tsc2^×/×^* ELT3 rat uterine leiomyoma cells, and *Tsc2^×/×^* mouse embryonic fibroblasts (MEFs) were treated with mizoribine for 48 hrs, alone and in combination with inhibitors of ATR, CHK1, CHK1/CHK2, WEE1, ATM, or DNA-PK added for the last 6-12 hrs. Inhibition of ATR, CHK1, and WEE1 enhanced IMPDH inhibitor-induced Caspase-3 cleavage in all three cell types, whereas the effects of ATM and DNA-PK inhibition appeared somewhat less robust and/or cell type-specific (Figure 2A-C), consistent with their reduced efficacy in preventing colony formation and regrowth (Supplemental Figure 1). As expected, ATR inhibition reduced phosphorylation of its direct substrate CHK1 Ser345, CHK1 inhibition reduced autophosphorylation on Ser296, and WEE1 inhibition reduced phosphorylation of its direct substrate CDK1 Tyr15 (Figure 2A-C). We were unable to measure the efficacy of ATM and DNA-PK inhibition in these cells due to well-known difficulties in acquiring antibodies that recognize activating phosphorylation of ATM (Ser1981) or DNA-PK (Ser2056), or phosphorylation of the direct ATM substrate CHK2 Thr68 in mouse/rat cells.

**Figure 2.**
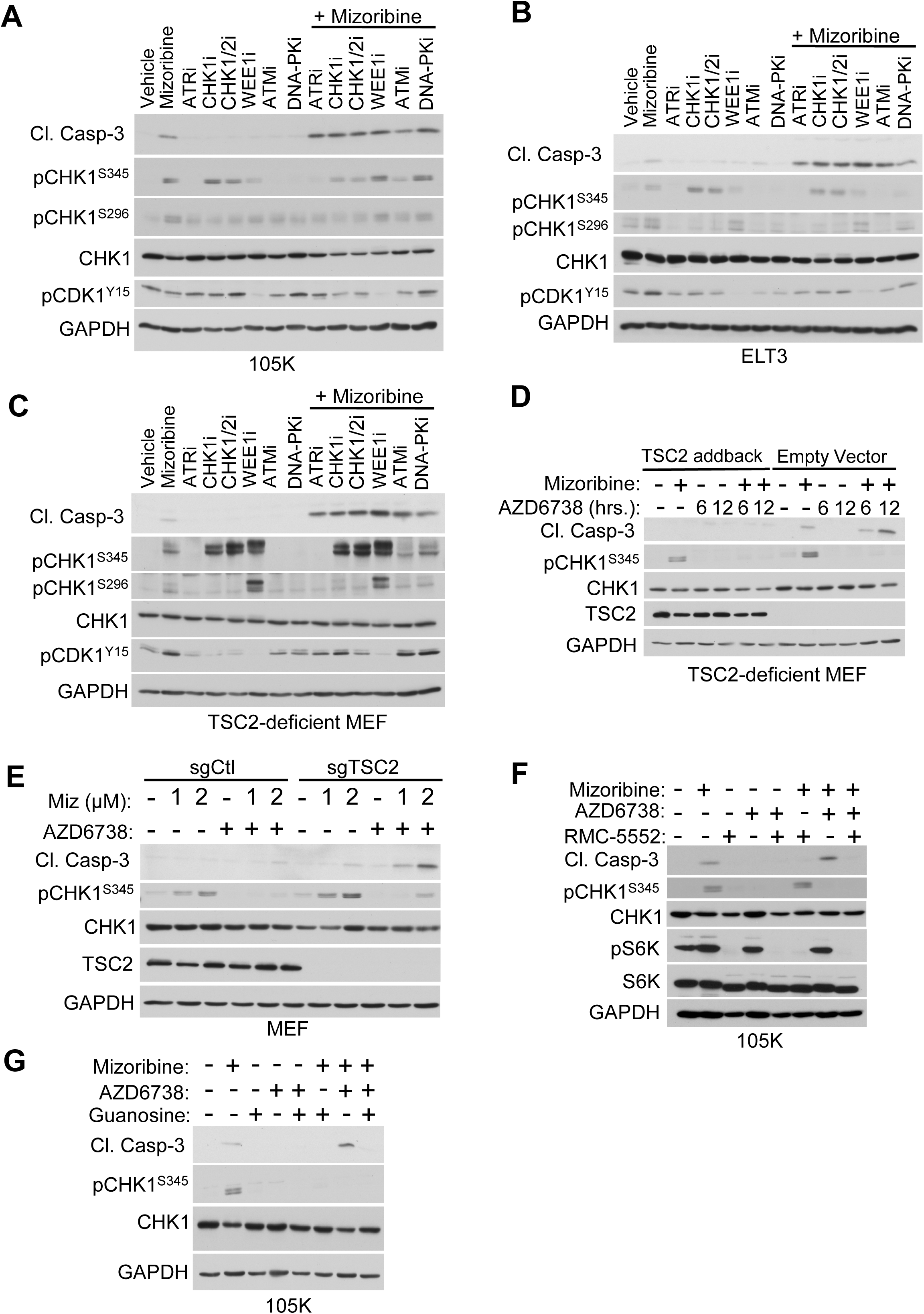
Inhibition of RSR/DDR kinases enhances IMPDH inhibitor-induced apoptosis while maintaining selectivity for TSC2-deficient cells. **(A)** *Tsc2^−/−^*105K renal tumor cells treated with mizoribine (3 µM) for 48 hrs, alone and in combination with indicated inhibitors added for the last 12 hrs, followed by immunoblotting for the indicated proteins. ATRi (AZD6738, 2 µM); CHK1i (LY2603618, 0.5 µM); Dual CHK1/2i (AZD7762, 0.125 µM); WEE1i (AZD1775, 0.125 µM); ATMi (AZD0156, 3 µM); DNA-PKi (AZD7648, 3 µM). **(B)** *Tsc2^−/−^* ELT3 rat uterine leiomyoma cells treated with mizoribine (5 µM) for 48 hrs, alone and in combination with indicated inhibitors added for the last 12 hrs, followed by immunoblotting for the indicated proteins. ATRi (AZD6738, 2 µM); CHK1i (LY2603618, 0.5 µM); Dual CHK1/2i (AZD7762, 0.125 µM); WEE1i (AZD1775, 0.125 µM); ATMi (AZD0156, 1 µM); DNA-PKi (AZD7648, 2 µM). **(C)** *Tsc2^−/−^ Trp53^−/−^* mouse embryonic fibroblasts (MEFs) treated with mizoribine (3 µM) for 48 hrs, alone and in combination with indicated inhibitors added for the last 6 hrs, followed by immunoblotting for the indicated proteins. ATRi (AZD6738, 2 µM); CHK1i (LY2603618, 0.5 µM); Dual CHK1/2i (AZD7762, 1 µM); WEE1i (AZD1775, 0.5 µM); ATMi (AZD0156, 1 µM); DNA-PKi (AZD7648, 2 µM). **(D)** *Tsc2^−/−^ Trp53^+/+^* 3T3 MEFs stably re-expressing TSC2 or empty vector control, treated with mizoribine (2 µM) for 48 hrs, alone and in combination with the ATR inhibitor AZD6738 (2 µM) added for the last 6 hrs or 12 hrs, as indicated, followed by immunoblotting for the indicated proteins. **(E)** Wild-type MEFs with CRISPR-mediated deletion of TSC2 (sgTSC2) or control (sgCtl) treated with indicated doses of mizoribine for 24 hrs, alone and in combination with the ATR inhibitor AZD6738 (2 µM) added for the last 6 hrs, followed by immunoblotting for the indicated proteins. **(F)** Cells in (A) treated with mizoribine (3 µM) and/or the mTORC1 inhibitor RMC-5552 (1 nM) for 48 hrs. The ATR inhibitor AZD6738 (2 µM) was added as indicated for the last 12 hrs, followed by immunoblotting for the indicated proteins. **(G)** Cells in (A) treated for 48 hrs with mizoribine (3 µM) with or without addition of exogenous guanosine (50 µM). The ATR inhibitor AZD6738 (2 µM) was added as indicated for the last 12 hrs, followed by immunoblotting for the indicated proteins.

Importantly, the selectivity for targeting TSC2-deficient cells was maintained with the combination treatments, which did not induce Caspase-3 cleavage in TSC2 addback 3T3 MEFs (*Tsc2^×/×^*cells stably re-expressing TSC2), or in control TSC2-expressing MEFs (sgCtl) compared to those with CRISPR-mediated *Tsc2* deletion (sgTSC2) (Figure 2D,E; Supplemental Figure 2A-C). Sustained mTORC1 signaling is required for sensitivity to IMPDH inhibition in TSC2-deficient cells (15, 16). Thus, we tested whether mTORC1 is required for efficacy of the combination treatments. Treatment with the bi-steric mTORC1 inhibitor RMC-5552 (40), or rapamycin, prevented Caspase-3 cleavage induced by combined inhibition of IMPDH and ATR, CHK1, or WEE1 in 105K and ELT3 cells (Figure 2F; Supplemental Figure 2D-I). Effects of the combination treatments were also fully rescued by supplementation with exogenous guanosine (Figure 2G; Supplemental Figure 2, J-L), which replenishes guanylate nucleotide pools through the purine salvage pathway, independent of IMPDH (15, 16, 41). Together these results demonstrate that the combination treatments further exploit the previously characterized mTORC1-dependent metabolic vulnerability that is triggered by guanylate depletion upon IMPDH inhibition.

Efficacy of the recently discovered IMPDH inhibitor gliocidin (31) was similarly enhanced by combined inhibition of ATR, CHK1, or WEE1 in TSC2-deficient cells, as measured by Caspase-3 cleavage, cleavage of the Caspase-3 substrate PARP, and Annexin V/Propidium Iodide (AV/PI) staining (Figure 3A-D, Supplemental Figure 3A). We next sought to determine whether this therapeutic strategy could be applicable in human cancer cells. We recently found that PTEN-loss in T-cell acute lymphoblastic leukemia (T-ALL) confers sensitivity to IMPDH inhibition through activation of AKT/mTORC1 signaling (17). The ATR inhibitor AZD6738 strongly enhanced the cell killing effects of gliocidin, as well as the FDA-approved IMPDH inhibitor mycophenolic acid (MPA), in PTEN-deficient Jurkat and CCRF-CEM human T-ALL cells (Figure 3E,F; Supplemental Figure 3B). Together, these results indicate that ATR-CHK1-WEE1 inhibition broadly potentiates the pro-apoptotic activity of structurally distinct IMPDH inhibitors across diverse cell models of mTORC1-driven growth.

**Figure 3.**
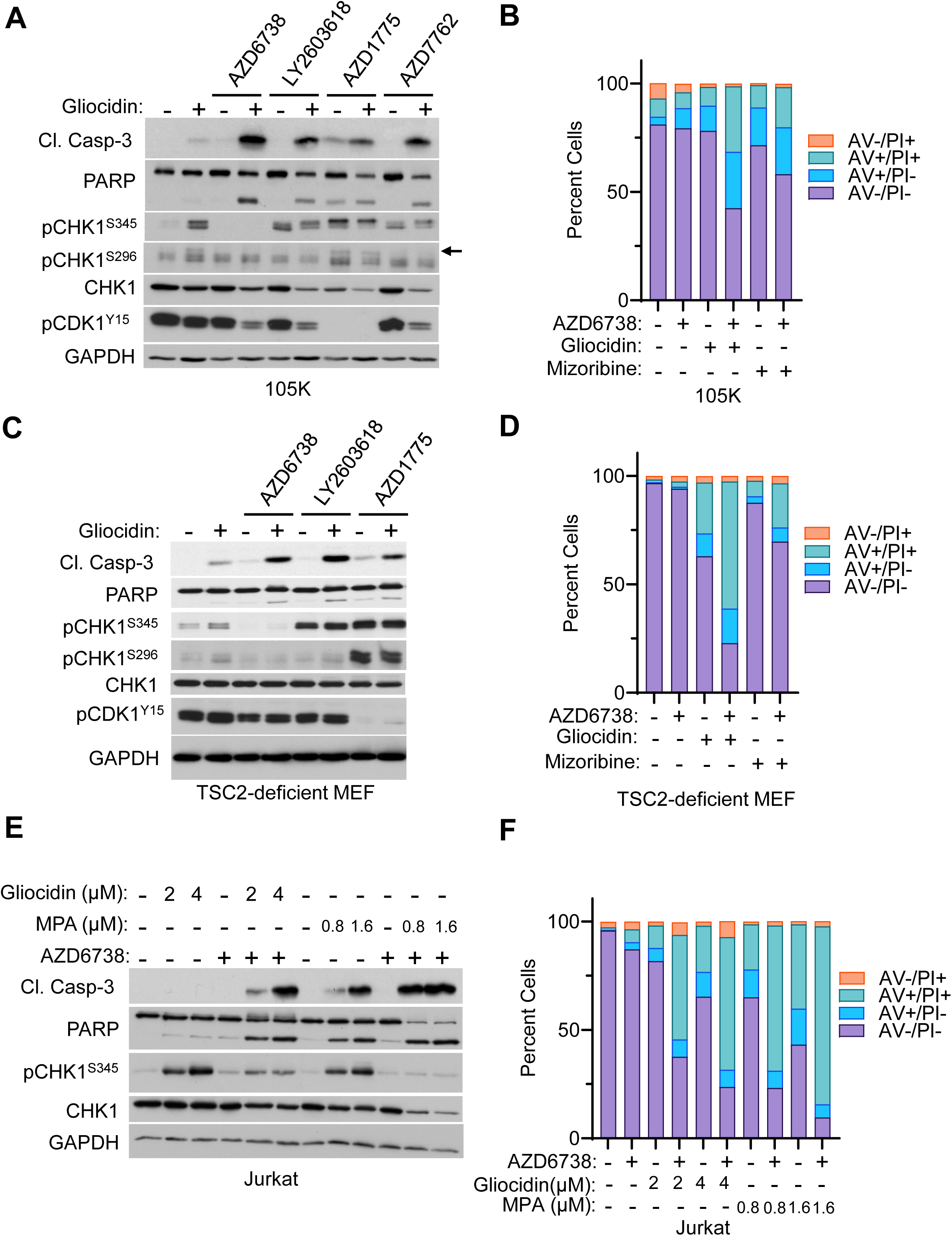
Inhibition of RSR/DDR kinases enhances apoptosis induced by structurally distinct IMPDH inhibitors in diverse cell types. **(A)** *Tsc2^−/−^* 105K renal tumor cells treated with the IMPDH inhibitor gliocidin (30 µM) for 48 hrs, alone and in combination with the ATR inhibitor AZD6738 (2 µM), CHK1 inhibitor LY2603618 (0.5 µM), WEE1 inhibitor AZD1775 (0.5 µM), or Dual CHK1/2 inhibitor AZD7762 (0.125 µM) added for the last 24 hrs, followed by immunoblotting for indicated proteins. **(B)** Cells in (A) treated with gliocidin (30 µM) or mizoribine (3 µM) for 72 hrs, alone and in combination with the ATR inhibitor AZD6738 (2 µM) added for the last 48hrs. Cell death was quantified by Annexin V/PI staining and graphed as percentage of the total cell population. **(C)** *Tsc2^−/−^ Trp53^−/−^* MEFs treated with gliocidin (2 µM) for 48 hrs, alone and in combination with the ATR inhibitor AZD6738 (2 µM), CHK1 inhibitor LY2603618 (0.5 µM), or WEE1 inhibitor AZD1775 (0.5 µM) added for the last 24 hrs, followed by immunoblotting for the indicated proteins. **(D)** Cells in (C) treated with gliocidin (4 µM) or mizoribine (3 µM) for 72 hrs, alone and in combination with the ATR inhibitor AZD6738 (2 µM) added for the last 48 hrs. Cell death was quantified by Annexin V/PI staining and graphed as percentage of the total cell population. **(E)** Jurkat T-cell acute lymphoblastic leukemia cells treated for 48 hrs with indicated doses of gliocidin or the FDA-approved IMPDH inhibitor mycophenolic acid (MPA), alone and in combination with the ATR inhibitor AZD6738 (1 µM), followed by immunoblotting for the indicated proteins. **(F)** Cells in (E) treated for 72 hrs with gliocidin or MPA, alone and in combination with the ATR inhibitor AZD6738 (1 µM) added for the last 48 hrs. Cell death was quantified by Annexin V/PI staining and graphed as percentage of the total cell population.

### ATR inhibition improves the durability of response to IMPDH inhibition in a syngeneic *Tsc2^−/−^* xenograft tumor model

We next tested the efficacy of combined inhibition of IMPDH and ATR *in vivo* in an immunocompetent, syngeneic xenograft model by establishing 105K xenograft tumors in C57BL/6J mice (Figure 4A) (16, 42, 43). Mice were treated for 30 days with vehicle, mizoribine (75 mg/kg/day by i.p. injection), AZD6738 (50 mg/kg Mon-Fri by oral gavage), or mizoribine plus AZD6738. The combination treatment was well tolerated, with no signs of toxicity or weight loss observed in any groups (Figure 4B). Mizoribine as a single agent exerts strong initial efficacy in this model (Figure 4C) (16). Nevertheless, addition of AZD6738 in combination with mizoribine reduced tumor volume by 40% on average compared to mizoribine alone (Figure 4C). To evaluate the durability of this response, treatments were discontinued after 30 days, and tumors were allowed to regrow for up to 6 months. Individual mice were euthanized as each tumor reached endpoint size (Figure 4D). Post-treatment survival was significantly extended in mice previously treated with the combination of mizoribine and AZD6738 compared to mizoribine monotherapy (Figure 4D). These findings demonstrate that ATR inhibition enhances the efficacy and durability of IMPDH inhibition *in vivo* in mTORC1-driven tumors.

**Figure 4.**
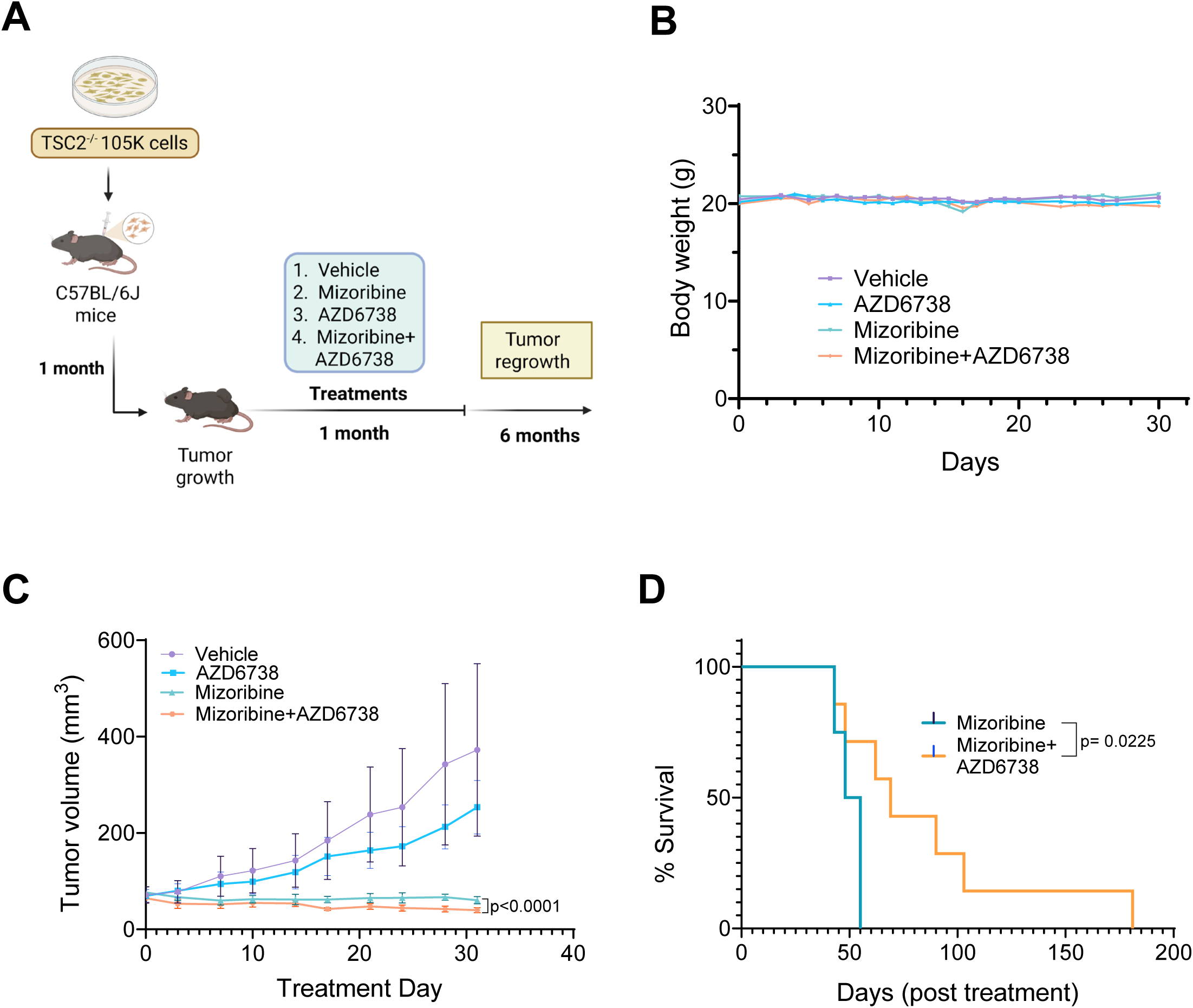
ATR inhibition improves the durability of response to IMPDH inhibition in a mouse *Tsc2^−/−^* xenograft tumor model. **(A)** Experimental design used in (B-D). C57BL/6J mice bearing syngeneic *Tsc2^−/−^* 105K xenograft tumors were treated for 30 days with vehicle, mizoribine (75 mg/kg/day by intraperitoneal injection), AZD6738 (50 mg/kg Mon-Fri by oral gavage), or mizoribine plus AZD6738. On Day 31, treatment was discontinued and tumors in the mizoribine and mizoribine plus AZD6738 groups were allowed to regrow until they reached endpoint. n = 7-8 mice per group. **(B)** Total body weight of the mice measured every day during the treatment phase, graphed as the mean of each group. **(C)** Tumor volume during the treatment phase measured every third day, graphed as mean ± SEM. *p < 0.0001 by two-way ANOVA. **(D)** Kaplan-Meier survival plots during the regrowth phase. p-value by Mantel-Cox log-rank test.

### ATR inhibition in combination with IMPDH inhibition causes aberrant cell cycle progression and micronuclei formation

To determine the mechanistic consequences of combined inhibition of IMPDH and ATR, we examined effects on underlying cell cycle dynamics. Mizoribine causes accumulation of Cyclin E1, accompanied by activating phosphorylation of CHK1 Ser345, and reduced DNA synthesis measured by 5-Ethynyl-2’-deoxyuridine (EdU) incorporation (Figure 5A-C; Supplemental Figure 4A,B). These effects are consistent with S-phase arrest and activation of the intra-S-phase checkpoint in response to nucleotide depletion (44), as previously reported in these cells (15, 16). Mizoribine also causes a modest increase in the levels of Cyclin B1 (Figure 5A,B), which begins to accumulate in S-phase and decreases sharply late in mitosis (45). In cells pre-treated with mizoribine, ATR inhibition rapidly reduced Cyclin E1 levels, indicating forced progression through S-phase, and later reduced Cyclin B1 levels, indicating that cells continue to progress through mitosis (Figure 5A,B). These trends continued with longer ATR inhibition (Supplemental Figure 4A,B), and similar effects were observed with inhibition of CHK1 or WEE1 (Supplemental Figure 4C,D). Arresting cells early in mitosis with nocodazole, the Eg5 inhibitor STLC (46), or at the G2/M checkpoint with the CDK1 inhibitor Ro-3306 (47) prevented the reduction in Cyclin B1 levels, confirming that ATR inhibition forces cell cycle progression from S-phase through mitosis (Figure 5B, Supplemental Figure 4E).

**Figure 5.**
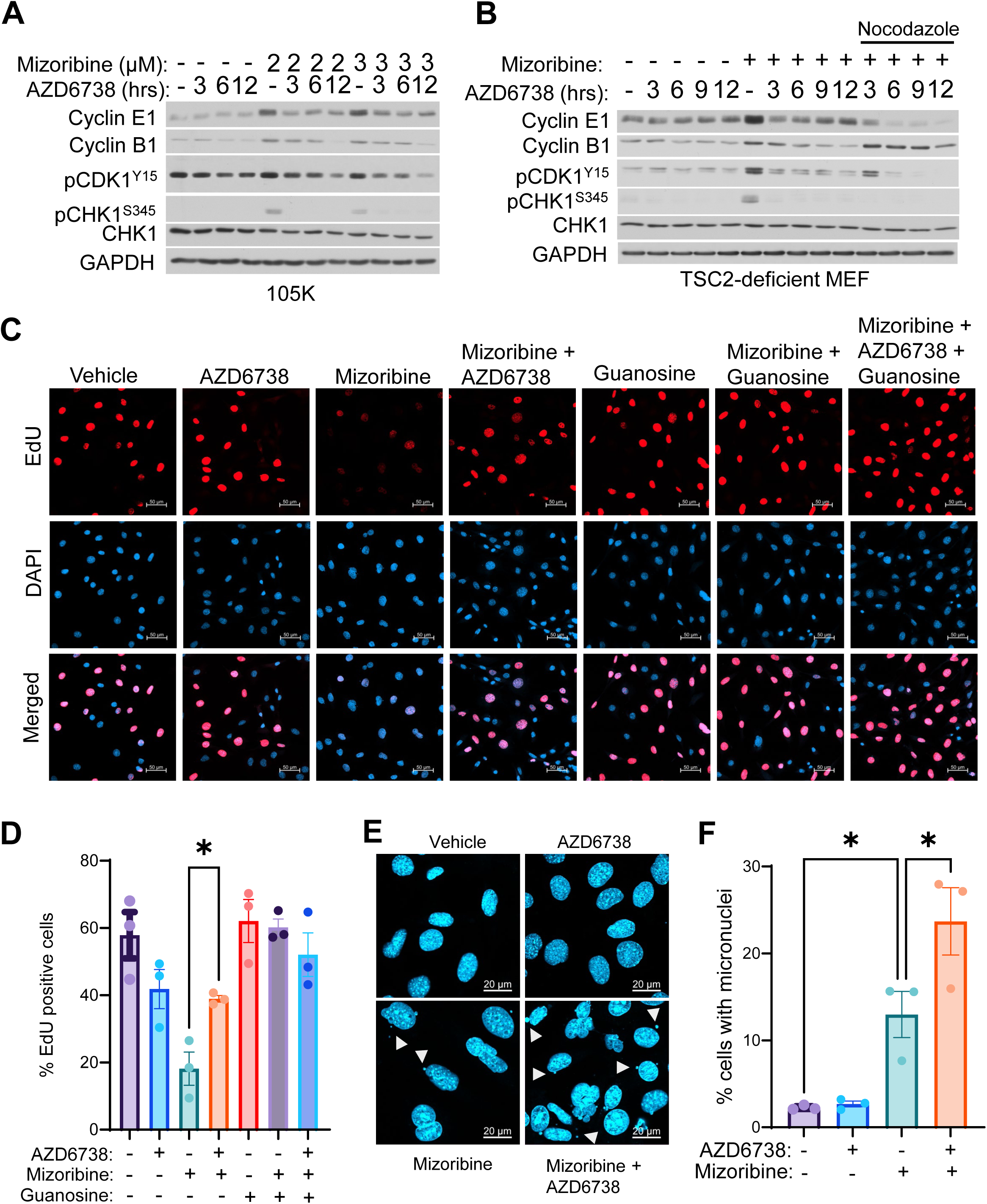
ATR inhibition in combination with IMPDH inhibition causes aberrant cell cycle progression and micronuclei formation. **(A)** *Tsc2^−/−^* 105K renal tumor cells treated with indicated doses of the IMPDH inhibitor mizoribine for 48 hrs, alone and in combination with the ATR inhibitor AZD6738 (2 µM) added for the last 3, 6, or 12 hrs as indicated, followed by immunoblotting for the indicated proteins. **(B)** *Tsc2^−/−^ Trp53^−/−^* MEFs treated with mizoribine (2 µM) for 48 hrs, alone and in combination with nocodazole (165 nM) and/or the ATR inhibitor AZD6738 (2 µM) added for the last 3, 6, 9, or 12 hrs as indicated, followed by immunoblotting for indicated proteins. **(C, D)** Cells in (B) treated for 24 hrs with mizoribine (2 µM) with or without supplementation with exogenous guanosine (50 µM). The ATR inhibitor AZD6738 (2 µM) was added as indicated for the last 6 hrs. Cells were pulse labeled with EdU (10 µM) for the last two hours, followed by DAPI staining. **(C)** Representative images. Scale bar = 50 µm. **(D)** EdU-positive cells were quantified and graphed as percentage of the total cell population. **(E, F)** Cells in (A) treated with mizoribine (3 µM) for 48 hrs, alone and in combination with the ATR inhibitor AZD6738 (2 µM) added for the last 12 hrs, followed by DAPI staining. **(E)** Representative images. White arrows indicate micronuclei. Scale bar = 20 µm. **(F)** Cells containing micronuclei were quantified and graphed as percentage of the total cell population. Graphical data are presented as mean of three biological replicates ± SEM. *p<0.05 by Welch’s t-test in (D) and one-way ANOVA followed by Dunnett’s multiple-comparison test in (F).

Interestingly, ATR inhibition partially rescued EdU incorporation in mizoribine-treated cells, indicating partial resumption of DNA synthesis as cell cycle checkpoints are abrogated (Figure 5C,D, Supplemental Figure 4F,G). This may be possible because IMPDH inhibition results in activation of the intra-S-phase checkpoint before guanylate nucleotide pools are completely exhausted, allowing cells to conserve a small guanylate pool (15–17), which could potentially be mobilized upon ATR inhibition. Supplementation with exogenous guanosine fully rescued EdU incorporation, demonstrating that guanylate pools are indeed limiting in cells treated with mizoribine or the combination of mizoribine plus AZD6738 (Figure 5C,D). Importantly, addition of AZD6738 to mizoribine-treated cells increased formation of micronuclei (Figure 5E,F), which are aberrant structures that contribute to genomic instability, forming when cells enter mitosis with damaged or under-replicated DNA (48). Together these data indicate that ATR inhibition abolishes the protective cell cycle checkpoints that limit IMPDH inhibitor efficacy in TSC2-deficient cells, forcing aberrant cell cycle progression and exacerbating cellular stress.

### Inhibition of ATR-CHK1-WEE1 signaling exacerbates IMPDH inhibitor-induced DNA damage

To determine whether the combination treatments induce DNA damage in TSC2-deficient cells, we measured the relative immunofluorescence staining intensity of the DNA double-strand break marker phospho-Histone H2AX Ser139 (p-H2AX) (49). Combining mizoribine with inhibition of ATR, CHK1 or WEE1 significantly increased p-H2AX staining intensity (Figure 6A,B, Supplemental Figure 5A,B). To directly measure DNA strand breakage, we performed alkaline comet assays (50). Treatment with mizoribine resulted in DNA tailing, which was significantly increased by combined inhibition of ATR (Figure 6C-F). Furthermore, immunofluorescence analysis of the DNA double-strand break repair factor 53BP1, which moves from diffuse nuclear staining to distinct puncta as it accumulates at the sites of DNA double-strand breaks (51), demonstrated that combining mizoribine with ATR inhibition increased the percent of cells harboring nuclear 53BP1 puncta (>5 puncta/cell) compared to mizoribine alone (Figure 6G,H). These results indicate that inhibition of ATR-CHK1-WEE1 signaling exacerbates IMPDH inhibitor-induced DNA damage.

**Figure 6.**
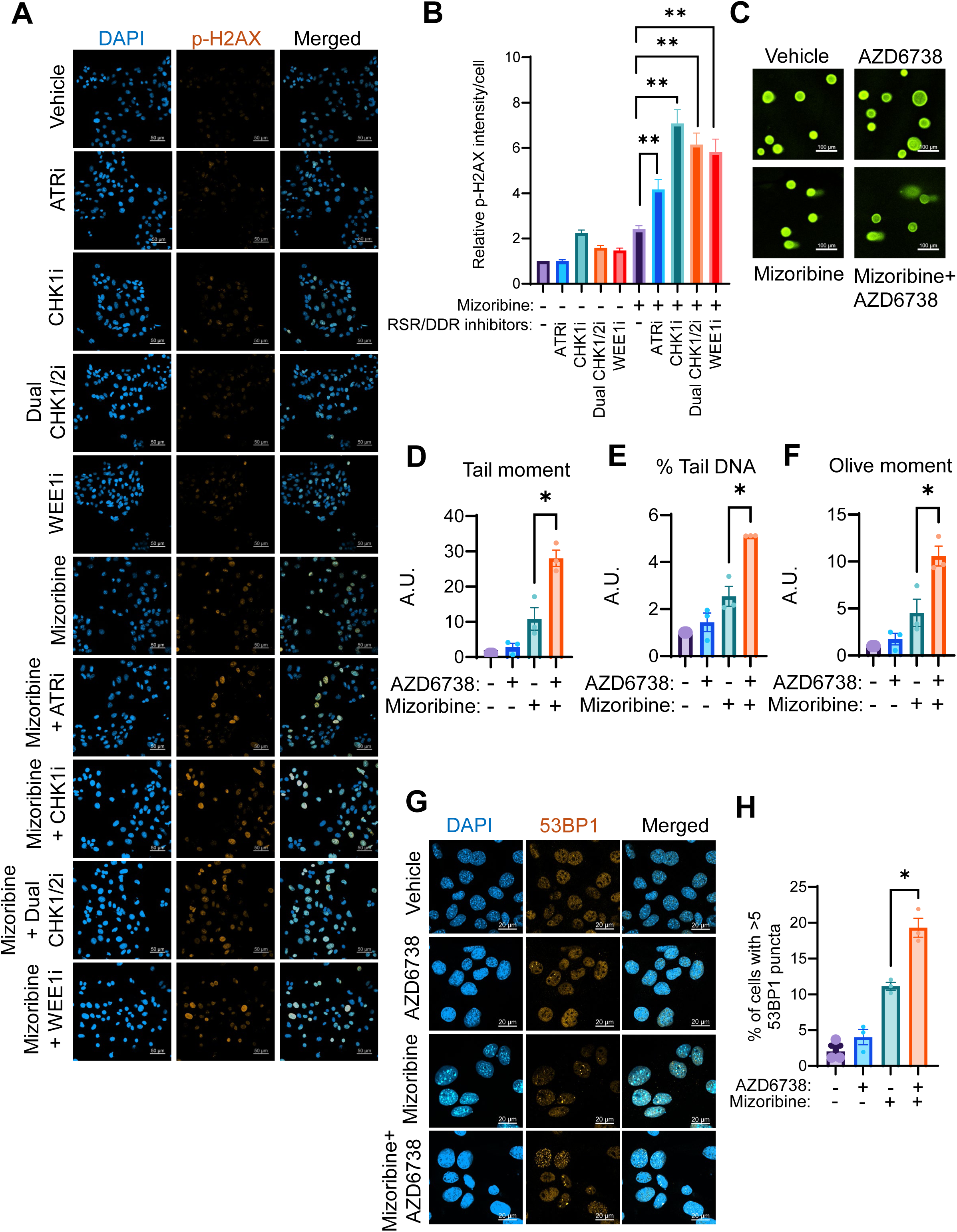
Inhibition of ATR-CHK1-WEE1 signaling exacerbates IMPDH inhibitor-induced DNA damage. **(A, B)** *Tsc2^−/−^* 105K renal tumor cells treated with mizoribine (2 µM) for 48 hrs, alone and in combination with indicated inhibitors added for the last 12 hrs, followed by immunofluorescence staining for p-H2AX S139 (p-H2AX). ATRi (AZD6738, 2 µM); CHK1i (LY2603618, 0.5µM); Dual CHK1/2i (AZD7762, 0.125µM); WEE1i (AZD1775, 0.125µM). **(A)** Representative images, scale bar = 50 µm. **(B)** p-H2AX staining intensity per cell was quantified and graphed relative to control. n = 150 cells per condition. **(C-F)** Alkaline Comet Assay on cells in (A) treated with mizoribine (10 µM) for 48 hrs, alone and in combination with the ATR inhibitor AZD6738 (2 µM) added for the last 12 hrs. **(C)** Representative images, scale bar = 100 µm. **(D)** Tail moment, **(E)** % Tail DNA, and **(F)** Olive tail moment were quantified and graphed as fold change relative to vehicle control. n = 20 cells per condition. **(G, H)** Cells in (A) treated with mizoribine (3 µM) for 48 hrs, alone and in combination with the ATR inhibitor AZD6738 (2 µM) added for the last 24 hrs, followed by immunofluorescence staining for 53BP1. **(G)** Representative images, scale bar = 20 µm. **(H)** 53BP1 puncta were quantified and graphed as percent of cells containing more than 5 puncta. Graphical data are presented as mean of three biological replicates ± SEM. *p<0.05, **p<0.001 by Welch’s t-test.

## Discussion

Here we find that inhibition of the ATR-CHK1-WEE1 signaling axis enhances the anti-tumor efficacy of IMPDH inhibitors in mTORC1-driven tumor cells. Combining the clinically approved IMPDH inhibitor mizoribine (currently used as an immunosuppressant in Asia) with clinically developed inhibitors of ATR (AZD6738/ceralasertib), CHK1 (LY2603618/rabusertib), WEE1 (AZD1775/adavosertib), or a dual CHK1/2 inhibitor (AZD7762), improves initial efficacy and slows regrowth when treatment is discontinued in cell models and *in vivo.* Critically, the combination treatments maintain robust selectivity for targeting mTORC1-driven tumor cells that has been previously characterized for IMPDH inhibitors as single agents (15–17). Correspondingly, the combination of mizoribine and AZD6738 significantly enhances efficacy and durability of response *in vivo,* compared to mizoribine alone, without signs of toxicity or weight loss (Figure 4). Given the immunosuppressive nature of IMPDH inhibitors, it is also notable that these effects are observed in immunocompetent, wild-type C57BL/6J mice (Figure 4).

Delayed cell and tumor regrowth upon cessation of the combination treatment corresponds to increased cell death (Figures 2,3), as well as the ability to induce cellular senescence, which does not occur with the single treatments (Figure 1D,E). The enhanced cytotoxic response also remains dependent on sustained mTORC1 signaling, as mTORC1 inhibition with RMC-5552 or rapamycin protects cells from apoptosis induced by the combination treatments (Figure 2F, Supplemental Figure 2D-I). Supplementation with exogenous guanosine fully rescues effects of the combination treatments, confirming that they stem from guanylate depletion (Figure 2G, 5C,D; Supplemental Figure 2J-K). Together these findings indicate a protective role for ATR-CHK1-WEE1 signaling in limiting IMPDH inhibitor efficacy. This protective role is abolished by the combination treatments, resulting in enhanced anti-tumor efficacy.

Inhibiting ATM or DNA-PK was less effective in combination with IMPDH inhibition (Supplemental Figure 1E-H, Figure 2A-C). ATR-CHK1-WEE1 mediate adaptive responses to DNA replication stress and DNA damage, whereas ATM and DNA-PK primarily respond to DNA damage (36–38, 52). IMPDH inhibition in mTORC1-driven tumor cells induces early DNA replication stress, with DNA damage accumulating later (15–17). Thus, the enhanced efficacy of IMPDH inhibition in combination with ATR-CHK1-WEE1 inhibition, compared to with ATM or DNA-PK inhibition, is consistent with an important role for blocking the early DNA replication stress response to enhance tumor cell-killing.

In addition to enhancing efficacy in TSC2-deficient cells and tumors, we also demonstrated that ATR inhibition enhances the cell-killing effects of IMPDH inhibitors in PTEN-deficient T-cell acute lymphoblastic leukemia cells (Figure 3E,F; Supplemental Figure 3B). We recently demonstrated that mTORC1 activation downstream of PTEN-loss sensitizes these cells to IMPDH inhibition (17). This indicates that the combination treatments could have potential for broad efficacy in diverse cancer settings driven by various mTORC1-activating mutations. Interestingly, PTEN deficiency was recently shown to sensitize high-grade serous ovarian cancer cells to ATR inhibition (53). IMPDH inhibition was also recently reported to induce DNA replication stress and sensitivity to ATR inhibition in Merkel cell carcinoma cells (54). Further studies are needed to uncover the full translational potential of this combination approach, including in PTEN-deficient cancers, as well as in small cell lung cancer, hepatocellular carcinoma, leukemia, and glioblastoma, where IMPDH inhibitors have strong efficacy as single agents in pre-clinical models (17, 22–31).

Mechanistically, our cell cycle analyses provide insight into how ATR-CHK1-WEE1 inhibition converts metabolic stress into acute cytotoxicity. IMPDH inhibition alone causes guanylate nucleotide depletion, activation of the intra-S-phase checkpoint, and S/G2 arrest (Figure 5, Refs (15–17). The intra-S-phase checkpoint, mediated by ATR-CHK1-WEE1, stabilizes stalled replication forks, prevents new replication origins from firing, and prevents cell cycle progression when DNA replication is incomplete (32–34). Inhibition of ATR, CHK1, or WEE1 abrogates this checkpoint, resulting in aberrant cell cycle progression in the presence of continued IMPDH inhibition, as measured by decreased levels of Cyclin E1 and Cyclin B1, and partial resumption of DNA synthesis (Figure 5, Supplemental Figure 4). This is accompanied by increased phospho-H2AX staining, nuclear 53BP1 foci, and comet assay tails, indicating increased DNA damage (Figure 6, Supplemental Figure 5). Entry into mitosis with damaged or under-replicated DNA leads to formation of micronuclei (Figure 5E,F). These data suggest that aberrant cell cycle progression during conditions of nucleotide depletion compromises genome integrity, potentially contributing to enhanced apoptosis and reduced proliferative recovery following combination treatment.

An important facet of this therapeutic strategy is the potential to repurpose clinically approved IMPDH inhibitors, including mizoribine and MPA/MMF, which have well-characterized pharmacokinetic and safety profiles from their use as immunosuppressants (20, 21, 55–59). In addition, the ATR, CHK1, and WEE1 inhibitors used in this study are undergoing clinical evaluation (36, 37, 60, 61). Our findings provide rationale for further investigation of combined IMPDH and ATR-CHK1-WEE1 inhibition in mTORC1-driven tumors, with the goal of enhancing tumor cell-killing and improving the durability of response compared to mTORC1 inhibitors. Given the prevalence of uncontrolled mTORC1 activation in cancer and genetic tumor syndromes, this combination approach has the potential for broad therapeutic applications.

## Methods

### Sex as a biological variable

Our study exclusively examined female mice. It is unknown whether the findings are relevant for male mice.

### Cell lines

*Tsc2*^×/×^ 105K mouse renal tumor-derived cells, *Tsc2*^×/×^ ELT3 rat uterine leiomyoma cells, *Tsc2*^×/×^; *Trp53^×/×^*primary mouse embryonic fibroblasts (MEFs), and *Tsc2*^×/×^; *Trp53*^+/+^ 3T3-immortalized MEFs stably reconstituted with wild-type TSC2 or empty vector were described previously (42, 62–64) and grown in DMEM (Corning #10-017-CV) supplemented with 10% heat-inactivated fetal bovine serum (ThermoFisher Scientific # A5256701) and 1% penicillin-streptomycin (Corning #30-002-Cl). To generate MEFs with CRISPR/Cas9-mediated *Tsc2* deletion (Figure 2E, Supplemental Figure 2B,C), MEFs stably expressing dominant-negative p53 (65) were provided by Dr. Piotr Sicinski and transfected with pSpCas9 BB-2A-Puro (PX459) V2.0 All-In-One plasmid vector (GenScript) expressing sgRNA targeting exon 3 of mouse *Tsc2* (ACAATCGCATCCGAATGATA), or a nontargeting control. Single-cell clones were grown in media supplemented with 2 μg/ml of puromycin and screened for loss of TSC2 expression and function by immunoblotting and immunofluorescence (Supplemental Figure 2B,C). Jurkat (DSMZ #ACC 282) and CCRF-CEM (DSMZ #ACC 240) cells were grown in RPMI-1640 (Corning #10-040-CV), supplemented with 10% heat-inactivated fetal bovine serum (ThermoFisher Scientific # A5256701) and 1% penicillin-streptomycin (Corning #30-002-Cl). Cells were maintained at 37°C in a humidified incubator with 5% CO₂. All cell lines were screened for mycoplasma using the Mycoplasma Detection Kit (SouthernBiotech #13100-01) every three months.

### Chemical compounds

The following compounds were resuspended in DMSO, unless otherwise indicated, and added into the cell culture medium at final concentrations indicated in the figure legends: mizoribine (Selleckchem #S1384) in water, mycophenolic acid (MPA, Selleckchem #S1501), gliocidin/SW106065 (MedChemExpress #HY-124778), AZD6738/ceralasertib (Selleckchem #S7693), LY2603618/rabusertib (Selleckchem #S2626), AZD7762 (Selleckchem #S1532), AZD1775/adavosertib (MedChemExpress #HY-10993), AZD0156 (Selleckchem #S8375), AZD7648 (Selleckchem #S8843), rapamycin (MedchemExpress #HY-10219), RMC-5552 (MedChemExpress #HY-132168) guanosine (Millipore-Sigma #G6752), nocodazole (Millipore-Sigma #M1404), S-trityl-L-cysteine (STLC) (MedChemExpress #HY-W011102), and Ro-3306 (MedChemExpress #HY-12529). Vehicle-treated cells received the corresponding drug solvents.

### Colony formation and regrowth assays

105K cells were seeded in 6-well plates at 1000 - 1200 cells/well and allowed to attach before treatment. Cells were treated for 6 days as indicated in the Figure Legends. At the end of treatment, cells were washed and stained with 0.5% (w/v) crystal violet to measure colony formation. In parallel plates, inhibitors were washed out after 6 days, and cells were allowed to regrow in fresh, drug-free, medium for an additional 3 days, followed by crystal violet staining. Plates were imaged using Azure Biosystems c600 Imager (AZ1600-01) and colony area was quantified using ImageJ.

### Cell viability assays

Cell viability was measured using the CellTiter-Glo Luminescent Cell Viability Assay (Promega G7573) according to the manufacturer’s instructions. Luminescence was measured using a SpectraMax iD3 Multimode Microplate Reader (Molecular Devices). Annexin V/PI staining was performed using the Dead Cell Apoptosis Kit (ThermoFisher Scientific #V13245), according to manufacturer’s instructions. Staining was measured with a Gallios Flow Cytometer (Beckman Coulter) and analyzed with Kaluza Analysis 2.1 software.

### Senescence-associated β-galactosidase staining

Cellular senescence was measured using the Senescence-Associated β-Galactosidase (SA-β-gal) staining kit (Cell Signaling Technology #9860) according to manufacturer’s instructions. SA-β-gal-positive cells were counted and graphed as percent of the total cell population.

### Immunoblotting and antibodies

Cells were lysed in in triton X-100 lysis buffer (20 mM Tris pH 7.5, 140 mM NaCl, 1 mM EDTA, 10% glycerol, 1% triton X-100, 1 mM DTT, 50 mM NaF) or RIPA buffer (150 mM NaCl, 1% IGEPAL, 0.5% sodium deoxycholate, 0.1%SDS, 50 mM Tris pH 8, 1 mM DTT), supplemented with protease inhibitor cocktail (Sigma #P8340), phosphate inhibitor cocktail #2 (Sigma #P5726) and #3 (Sigma #P0044) at 1:100 each. Western blotting was performed using the following antibodies at 1:1000 dilution: Cleaved caspase-3 (CST #9664), PARP (CST #9542), pCHK1^S345^ (CST #2348), pCHK1^S296^ (CST #90178), total CHK1 (CST #2360), pCDK1^Y15^ (CST #4539), Cyclin E1 (CST #20808), Cyclin B1 (CST #4138), pS6K^T389^ (CST #9234), total S6K (CST #2708), Raptor (CST #2280), TSC1 (CST #6935), TSC2 (CST #4308), and GAPDH (CST #5174).

### Immunostaining

Cells were grown on glass coverslips and fixed with 4% formaldehyde (ThermoFisher Scientific #28908) for 15 min at room temperature, then washed with PBS. Cells were blocked in PBS containing 5% normal goat serum (CST #5425) and 0.3% Triton X-100 for 60 min at room temperature, before incubation with primary antibodies TSC2 (CST #4308, 1:1250), p-H2AX^S139^ (CST #9718, 1:100), or 53BP1 (Novus Biologicals #NB100-304SS, 1:1000) overnight at 4°C. Cells were subsequently incubated with Cy3-conjugated secondary antibody (Jackson ImmunoResearch #111-165-144, 1:1000), for 1 hr at room temperature, counterstained with DAPI (0.5 μg/mL, ThermoFisher Scientific # D1306), and mounted with Fluoromount G (SouthernBiotech #0100-01). Images were acquired using a Zeiss LSM 900 confocal microscope and analyzed using Zeiss Zen Blue 3.3 software.

### EdU incorporation

Cells were grown on glass coverslips, treated as indicated in the Figure Legends, and pulse-labeled with 5-Ethynyl-2’-deoxyuridine (EdU, 10 μM, Lumiprobe #10540) for the final 2 hours of treatment to label newly synthesized DNA. Cells were fixed with 4% formaldehyde (ThermoFisher Scientific #28908) for 15 min at room temperature, then washed with PBS and permeabilized with 0.2% Tween-20 for 10 min at room temperature. Cells were blocked in PBS containing 3% BSA (Sigma, #A7030) for 1 hr at room temperature or overnight at 4°C. For EdU detection, Click-iT labeling was performed using 4 µM Sulfo-Cy5 azide (Lumiprobe #A3330) in PBS with 1 mM CuSO4 (freshly prepared) and 20 mg/mL Na-ascorbate (freshly prepared). Nuclei were counterstained with DAPI (0.5 μg/mL, ThermoFisher Scientific # D1306), and mounted with Fluoromount G (SouthernBiotech #0100-01). Images were acquired using a Zeiss LSM 900 confocal microscope, quantified, and graphed as percent of EdU positive cells.

### Mouse studies

All animal experiments were approved by the Rutgers Institutional Animal Care and Use Committee (IACUC). 105K xenograft tumors were established by injecting 1.2 million cells in a 1:1 mixture with Matrigel (Corning #356237) subcutaneously into the flank of syngeneic 6-week-old female C57BL/6J mice (Jackson Laboratory #000664). 7-8 mice per group were treated with vehicle, mizoribine (75 mg/kg/day in saline, by i.p. injection), AZD6738 (50 mg/kg Mon-Fri in saline +10% DMSO, 40% PEG-300, 5% Tween-80, by oral gavage), or the combination, beginning 4 weeks after cell injection when tumors became palpable. Mice were assigned to treatment groups based on their tumor size so that the average tumor size per group was equal among all groups at the start of treatment. Treatment continued for 30 days, with body weight measured daily, and tumor size measured every 3 days using digital calipers. Tumor volumes were calculated as (length x width^2^) / 2. At the end of 30-day treatment period, treatments were discontinued and tumors were allowed to regrow until reaching endpoint size, defined according to Rutgers IACUC as 20 mm length in any direction. Mice were continuously monitored throughout the experiment for signs of toxicity or weight loss.

### Comet assay

Comet assays were performed using the CometAssay Kit (Trevigen, # 4250-050-k). Cells were treated as indicated in the Figure Legends, trypsinized and embedded in agarose on glass slides, followed by lysis, alkaline DNA unwinding, and electrophoresis according to the manufacturer’s protocol. DNA was stained with SYBR Gold (ThermoFisher Scientific #S11494, 1:30,000 dilution in TE buffer (10 mM Tris pH 7.5, 1 mM EDTA)) and visualized using a Zeiss LSM 900 confocal microscope. Images were analyzed using Tritek software v2.0.0.38 for quantification of comet tail moment, percent of DNA in the tail, and olive tail moment.

### Statistics

Graphical data are presented as mean of at least 3 biological replicates ± SEM, unless otherwise indicated in the Figure Legends. Statistical analyses were performed using GraphPad Prism 11.01.0. Pairwise comparisons were determined by two-tailed Welch’s *t* test where indicated. One-way ANOVA followed by Dunnett’s multiple-comparison test was used for parametric comparison among multiple treatment groups. Two-way ANOVA followed by Bonferroni’s multiple comparisons test was used to compare treatment groups across different time points. Kaplan-Meier survival curves were analyzed by Mantel-Cox log-rank test. Statistical details for individual experiments can be found in their respective Figure Legends.

### Study approval

All animal experiments were approved by the Rutgers Institutional Animal Care and Use Committee (IACUC). Animal procedures were conducted in accordance with institutional guidelines for the care and use of laboratory animals.

### Data availability

Quantitative data from this study are available in the accompanying Supporting Data Values file, including numerical values used in all graphs in the main and supplemental figures. Additional experimental data and information are available from the corresponding author.

## Author contributions

Conceptualization (M.A., A.J.V.), conducting experiments (M.A., A.G., R.A.P., J.N.J., N.S., J.H.), analyzing data (M.A., A.J.V.), reagents and resources (S.F.B., A.J.V.), writing the manuscript (M.A., A.J.V.), manuscript review and editing (M.A., R.A.P., J.N.J., N.S., S.F.B., A.J.V.), project administration (A.J.V.).

## AI disclosure

The authors declare that no AI tools were used in the original research for, or writing of, this article.

## Acknowledgements

We thank Sumit Patel, Robert Teagan Polotaye, Daniel Herranz, Bing Xia, and Advaitha Madireddy for helpful discussions and technical assistance. We also thank Rutgers EOHSI (Environment and Occupational Health Sciences Institute) Flow Cytometry Core members Rita Hahn, Jessica Cervelli and Raymond Rancourt for technical assistance.

