## Supplemental Figures for "Targeting ATR-CHK1-WEE1 signaling enhances IMPDH inhibitor efficacy in mTORC1-driven tumor cells"

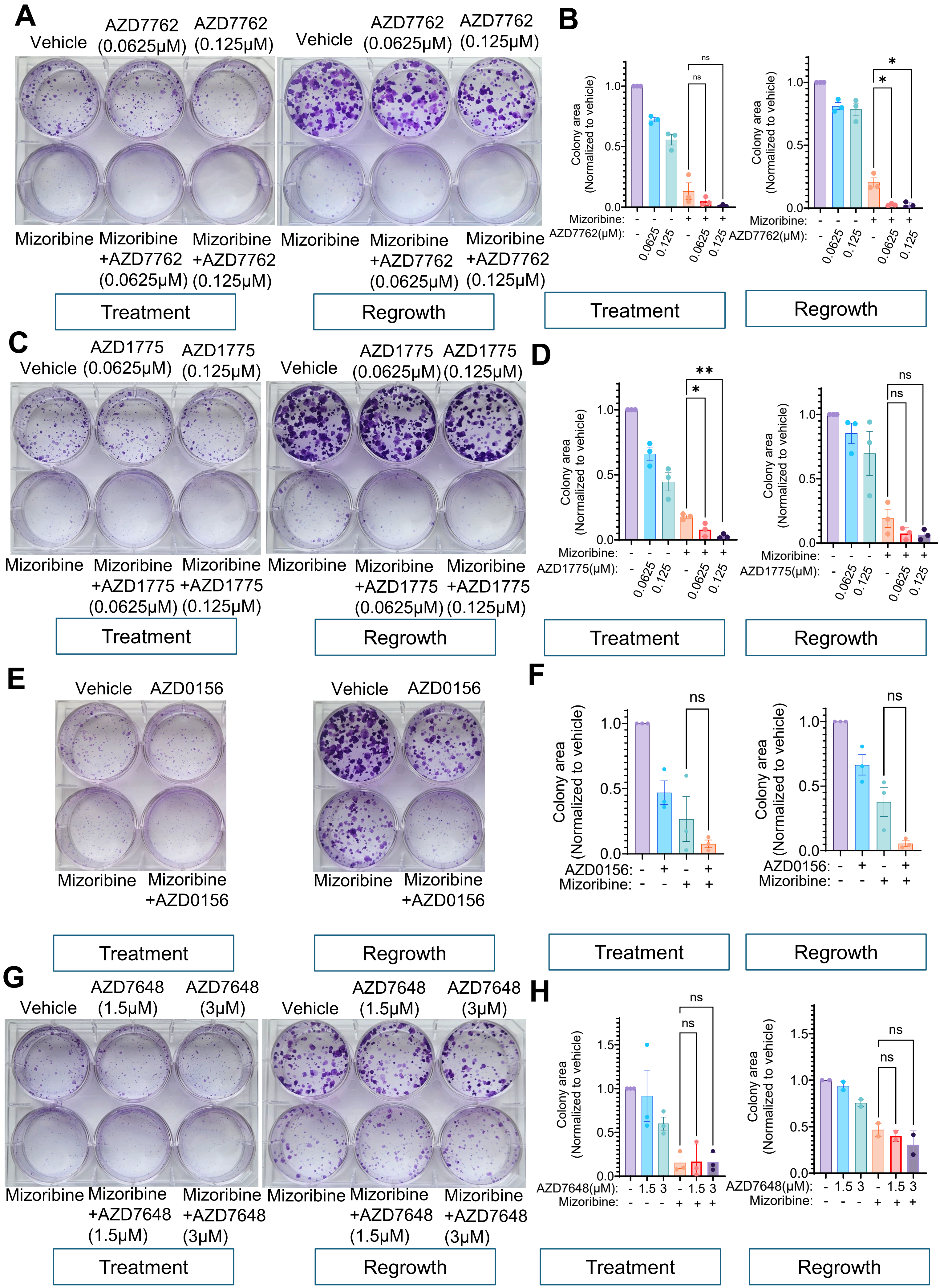

A
Vehicle
AZD7762 (0.0625μM)
AZD7762 (0.125μM)
Mizoribine
Mizoribine +AZD7762 (0.0625μM)
Mizoribine +AZD7762 (0.125μM)
Treatment
Regrowth
B
Colony area (Normalized to vehicle)
Mizoribine:
AZD7762(μM):
0.0625
0.125
ns
*
C
AZD1775 (0.0625μM)
AZD1775 (0.125μM)
Mizoribine +AZD1775 (0.0625μM)
Mizoribine +AZD1775 (0.125μM)
D
AZD1775(μM):
**
E
AZD0156
Mizoribine +AZD0156
F
AZD0156:
G
AZD7648 (1.5μM)
AZD7648 (3μM)
Mizoribine +AZD7648 (1.5μM)
Mizoribine +AZD7648 (3μM)
H
AZD7648(μM):

**Supplemental Figure 1. Supplemental data supporting Figure 1**

**(A, B)** *Tsc2*$^{-/-}$ 105K renal tumor cells were treated for 6 days with the IMPDH inhibitor mizoribine (3 μM) alone and in combination with indicated doses of the dual CHK1/2 inhibitor AZD7762. Following treatment, parallel plates were washed, replenished with fresh media, and allowed to regrow in the absence of drug treatments for an additional 3 days. **(A)** Colonies were visualized by crystal violet staining. **(B)** Colony area in (A) was quantified and graphed as fold change relative to the vehicle.

**(C-H)** Cells in (A) treated as described in (A) with mizoribine (3 μM) alone and in combination with indicated doses of the WEE1 inhibitor AZD1775 **(C, D)**, ATM inhibitor AZD0156 (30 nM) **(E, F)**, or DNA-PK inhibitor AZD7648 **(G, H)**. Colonies in (C, E, G) were visualized and quantified in (D, F, H), respectively, as in (A, B).

Graphical data are presented as mean of at least three biological replicates ± SEM. *$p<0.05$, **$p<0.001$ by Welch's t-test.

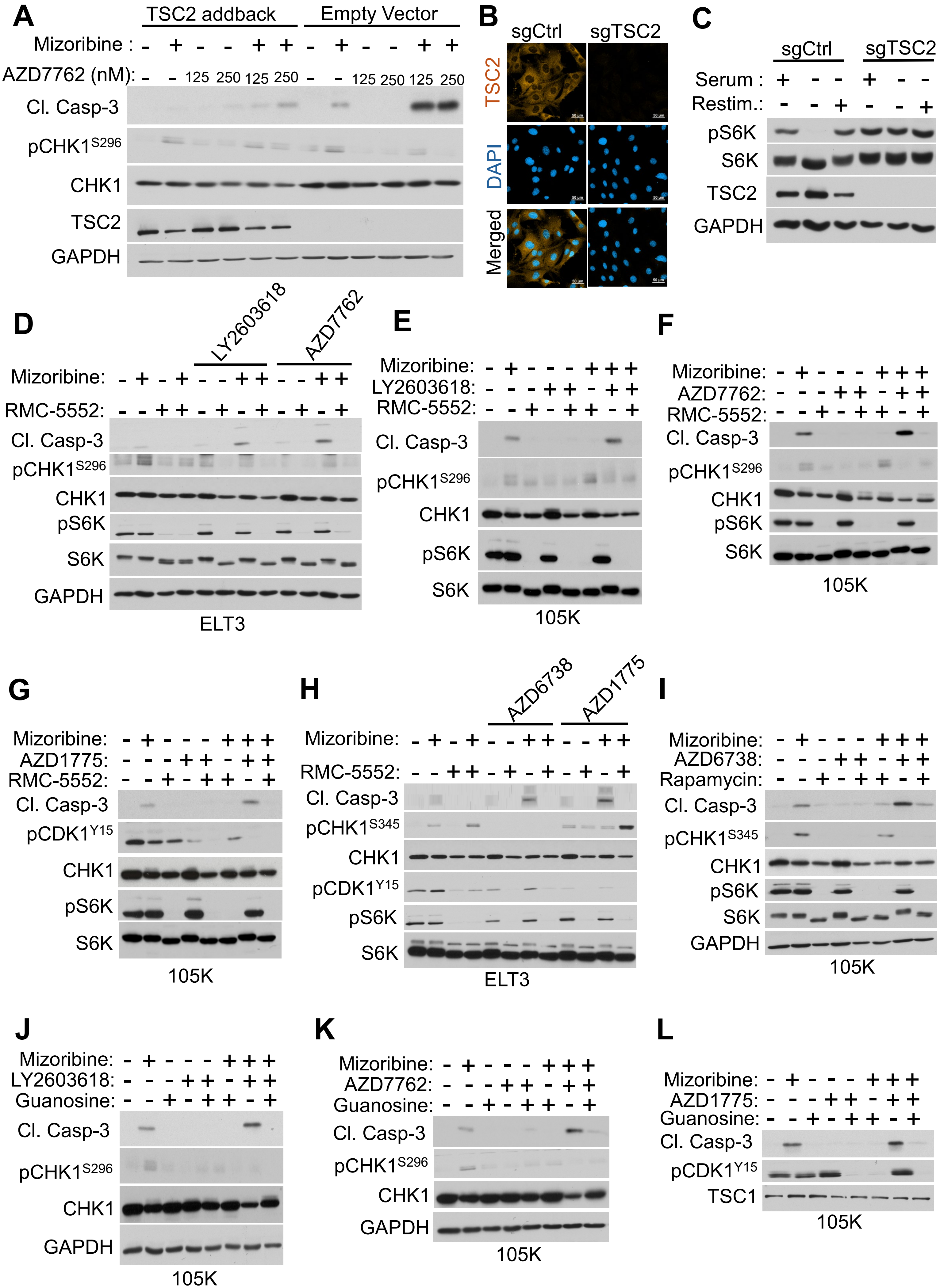
A
TSC2 addback
Empty Vector
Mizoribine : - + - - + + - + - - + +
AZD7762 (nM): - - 125 250 125 250 - - 125 250 125 250
Cl. Casp-3
pCHK1S296
CHK1
TSC2
GAPDH
B
sgCtrl
sgTSC2
TSC2
DAPI
Merged
C
sgCtrl
sgTSC2
Serum : + - - + - -
Restim.: - - + - - +
pS6K
S6K
TSC2
GAPDH
D
LY2603618
AZD7762
Mizoribine: - + - + - - + + - - + +
RMC-5552: - - + + - + - + - + - +
Cl. Casp-3
pCHK1S296
CHK1
pS6K
S6K
GAPDH
ELT3
E
Mizoribine: - + - - - + + +
LY2603618: - - - + + - + +
RMC-5552: - - + - + + - +
Cl. Casp-3
pCHK1S296
CHK1
pS6K
S6K
105K
F
Mizoribine: - + - - - + + +
AZD7762: - - - + + - + +
RMC-5552: - - + - + + - +
Cl. Casp-3
pCHK1S296
CHK1
pS6K
S6K
105K
G
Mizoribine: - + - - - + + +
AZD1775: - - - + + - + +
RMC-5552: - - + - + + - +
Cl. Casp-3
pCDK1Y15
CHK1
pS6K
S6K
105K
H
AZD6738
AZD1775
Mizoribine: - + - + - - + + - - + +
RMC-5552: - - + + - + - + - + - +
Cl. Casp-3
pCHK1S345
CHK1
pCDK1Y15
pS6K
S6K
ELT3
I
Mizoribine: - + - - - + + +
AZD6738: - - - + + - + +
Rapamycin: - - + - + + - +
Cl. Casp-3
pCHK1S345
CHK1
pS6K
S6K
GAPDH
105K
J
Mizoribine: - + - - - + + +
LY2603618: - - - + + - + +
Guanosine: - - + - + + - +
Cl. Casp-3
pCHK1S296
CHK1
GAPDH
105K
K
Mizoribine: - + - - - + + +
AZD7762: - - - + + - + +
Guanosine: - - + - + + - +
Cl. Casp-3
pCHK1S296
CHK1
GAPDH
105K
L
Mizoribine: - + - - - + + +
AZD1775: - - - + + - + +
Guanosine: - - + - + + - +
Cl. Casp-3
pCDK1Y15
TSC1
105K

**Supplemental Figure 2. Supplemental data supporting Figure 2**

**(A)** *Tsc2*$^{-/-}$ 3T3 MEFs stably re-expressing TSC2 or empty vector control, treated with mizoribine (2 μM) for 48 hrs, alone and in combination with indicated doses of the Dual CHK1/2 inhibitor AZD7762 added for the last 12 hrs, followed by immunoblotting for the indicated proteins.

**(B)** TSC2 immunofluorescence staining in wild-type MEFs with CRISPR-mediated deletion of TSC2 (sgTSC2) or control (sgCtl). Scale bar = 50 μm.

**(C)** Cells in (B) grown in complete media or serum starved for 24 hrs, with or without restimulation with serum for 15 min ("Restim").

**(D)** *Tsc2*$^{-/-}$ ELT3 rat uterine leiomyoma cells treated with mizoribine (5 μM) and/or the mTORC1 inhibitor RMC-5552 (1 nM) for 48 hrs. The CHK1 inhibitor LY2603618 (0.5 μM) or Dual CHK1/2 inhibitor AZD7762 (0.125 μM) were added as indicated for the last 12 hrs, followed by immunoblotting for the indicated proteins.

**(E-G)** *Tsc2*$^{-/-}$ 105K renal tumor cells treated with mizoribine (3 μM) and/or RMC-5552 (1 nM) for 48 hrs. **(E)** The CHK1 inhibitor LY2603618 (0.5 μM), **(F)** Dual CHK1/2 inhibitor AZD7762 (0.125 μM), or **(G)** WEE1 inhibitor AZD1775 (0.125 μM) were added for the last 12 hrs, followed by immunoblotting for the indicated proteins.

**(H)** Cells in (D) treated with mizoribine (5 μM) and/or RMC-5552 (1 nM) for 48 hrs. The ATR inhibitor AZD6738 (2 μM) or WEE1 inhibitor AZD1775 (0.125 μM) were added as indicated for the last 12 hrs, followed by immunoblotting for the indicated proteins.

**(I)** Cells in (E-G) treated with mizoribine (3 μM) and/or rapamycin (20 nM) for 48 hrs. The ATR inhibitor AZD6738 (2 μM) was added as indicated for the last 12 hrs, followed by immunoblotting for the indicated proteins.

**(J-L)** Cells in (E-G) treated for 48 hrs with mizoribine (3 μM), with or without addition of exogenous guanosine (50 μM). **(J)** The CHK1 inhibitor LY2603618 (0.5 μM), **(K)** Dual CHK1/2 inhibitor AZD7762 (0.125 μM), or **(L)** WEE1 inhibitor AZD1775 (0.125 μM) were added for the last 12 hrs, followed by immunoblotting for the indicated proteins.

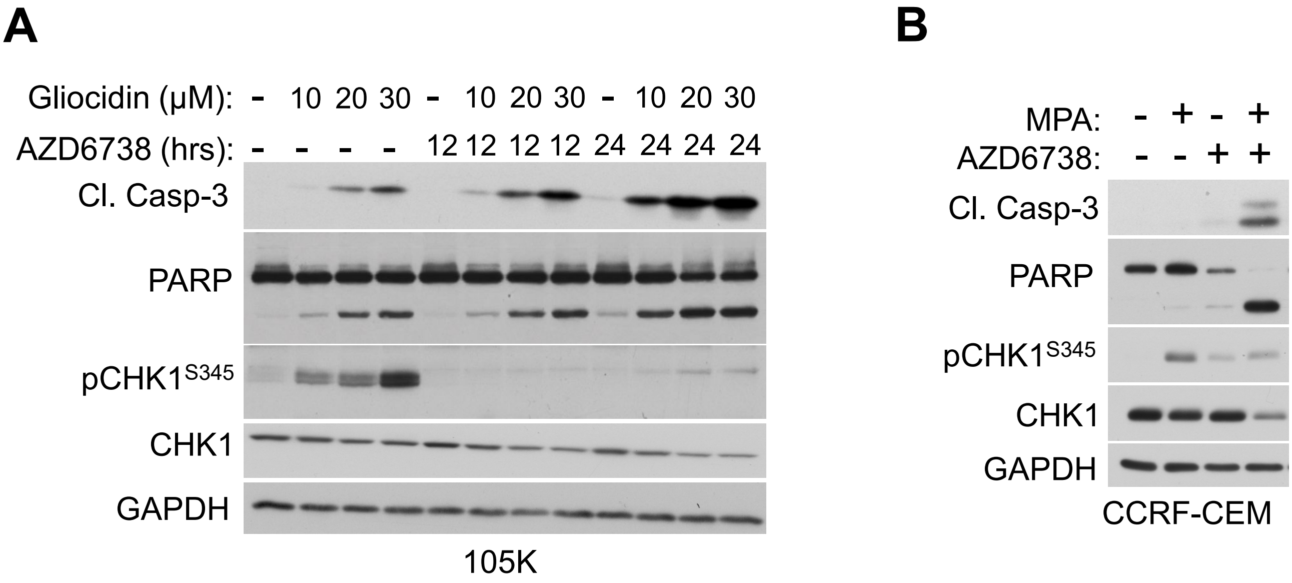
A
Gliocidin (μM): - 10 20 30 - 10 20 30 - 10 20 30
AZD6738 (hrs): - - - - 12 12 12 12 24 24 24 24
Cl. Casp-3
PARP
pCHK1S345
CHK1
GAPDH
105K
B
MPA: - + - +
AZD6738: - - + +
Cl. Casp-3
PARP
pCHK1S345
CHK1
GAPDH
CCRF-CEM

**Supplemental Figure 3. Supplemental data supporting Figure 3**

**(A)** *Tsc2*$^{-/-}$ 105K cells treated for 48 hrs with indicated doses of the IMPDH inhibitor gliocidin, alone and in combination with the ATR inhibitor AZD6738 (2 μM) added for the last 12 or 24 hrs, followed by immunoblotting for the indicated proteins.

**(B)** CCRF-CEM T-cell acute lymphoblastic leukemia cells treated with the FDA-approved IMPDH inhibitor mycophenolic acid (MPA, 1 μM) for 24 hrs, alone and in combination with the ATR inhibitor AZD6738 (2 μM) added for the last 12 hrs, followed by immunoblotting for the indicated proteins.

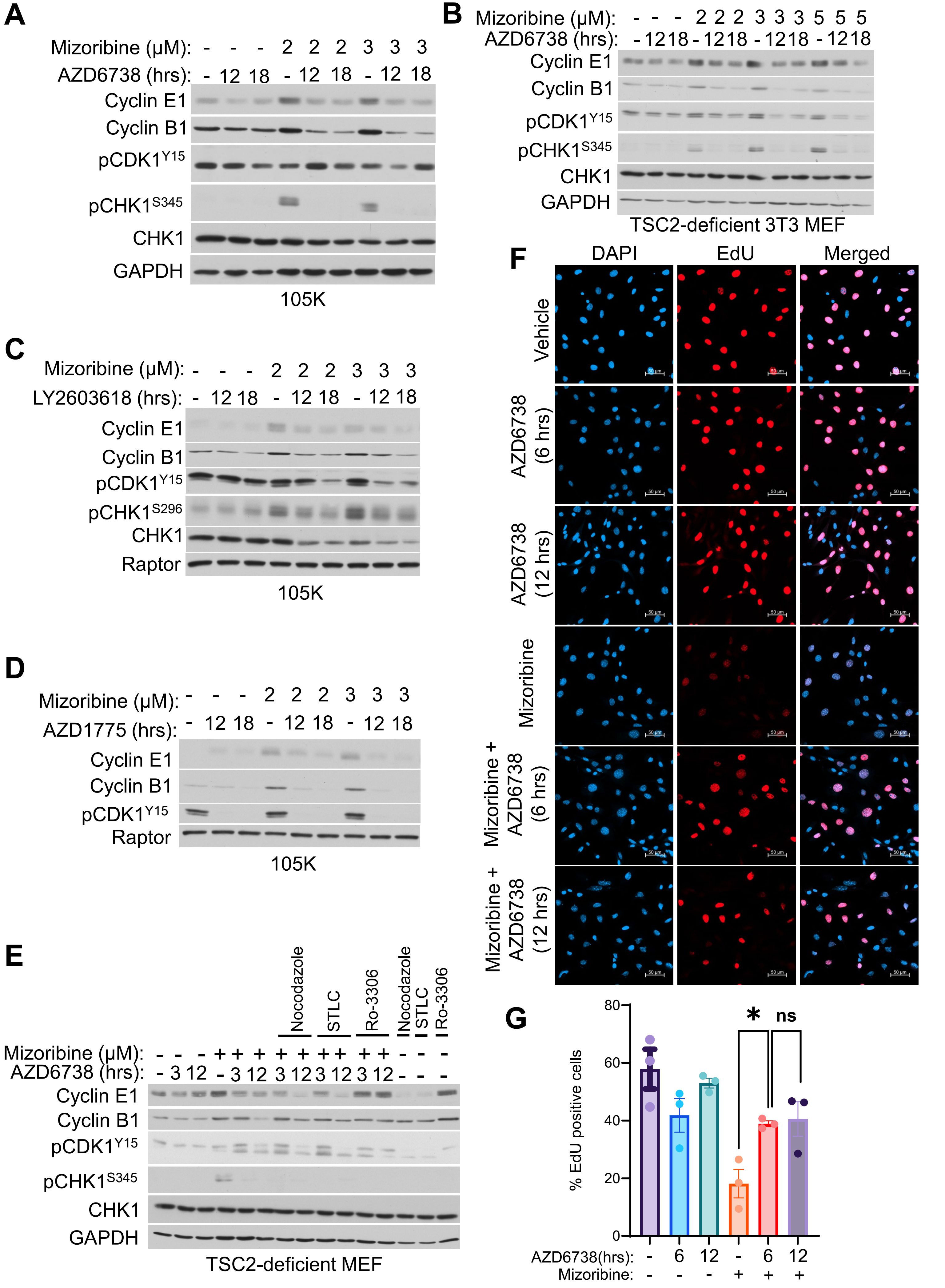
A
Mizoribine (µM): - - - 2 2 2 3 3 3
AZD6738 (hrs): - 12 18 - 12 18 - 12 18
Cyclin E1
Cyclin B1
pCDK1Y15
pCHK1S345
CHK1
GAPDH
105K
B
Mizoribine (µM): - - - 2 2 2 3 3 3 5 5 5
AZD6738 (hrs): - 12 18 - 12 18 - 12 18 - 12 18
Cyclin E1
Cyclin B1
pCDK1Y15
pCHK1S345
CHK1
GAPDH
TSC2-deficient 3T3 MEF
C
Mizoribine (µM): - - - 2 2 2 3 3 3
LY2603618 (hrs): - 12 18 - 12 18 - 12 18
Cyclin E1
Cyclin B1
pCDK1Y15
pCHK1S296
CHK1
Raptor
105K
D
Mizoribine (µM): - - - 2 2 2 3 3 3
AZD1775 (hrs): - 12 18 - 12 18 - 12 18
Cyclin E1
Cyclin B1
pCDK1Y15
Raptor
105K
E
Nocodazole
STLC
Ro-3306
Nocodazole
STLC
Ro-3306
Mizoribine (µM): - - - + + + + + + + + + - - -
AZD6738 (hrs): - 3 12 - 3 12 3 12 3 12 3 12 - - -
Cyclin E1
Cyclin B1
pCDK1Y15
pCHK1S345
CHK1
GAPDH
TSC2-deficient MEF
F
DAPI
EdU
Merged
Vehicle
AZD6738 (6 hrs)
AZD6738 (12 hrs)
Mizoribine
Mizoribine + AZD6738 (6 hrs)
Mizoribine + AZD6738 (12 hrs)
50 µm
G
*
ns
% EdU positive cells
80
60
40
20
0
AZD6738(hrs): - 6 12 - 6 12
Mizoribine: - - - + + +

**Supplemental Figure 4. Supplemental data supporting Figure 5**

**(A, B)** *Tsc2*$^{-/-}$ 105K renal tumor cells **(A)**, or *Tsc2*$^{-/-}$ 3T3 MEFs **(B)**, treated with indicated doses of the IMPDH inhibitor mizoribine for 48 hrs, alone and in combination with the ATR inhibitor AZD6738 (2 μM) added for the last 12 or 18 hrs as indicated, followed by immunoblotting for indicated proteins.

**(C, D)** Cells in (A) treated with indicated doses of mizoribine for 48 hrs, alone and in combination with the CHK1 inhibitor LY2603618 (0.5 μM) **(C)**, or WEE1 inhibitor AZD1775 (0.5 μM) **(D)**, added for the last 12 or 18 hrs as indicated, followed by immunoblotting for indicated proteins.

**(E)** *Tsc2*$^{-/-}$ *Trp53*$^{-/-}$ MEFs treated with mizoribine (2 μM) for 48 hrs, alone and in combination with nocodazole (165 nM), the Eg5 inhibitor STLC (10 μM), CDK1 inhibitor Ro-3306 (10 μM), and/or ATR inhibitor AZD6738 (2 μM) added for the last 3 or 12 hrs as indicated, followed by immunoblotting for indicated proteins.

**(F, G)** Cells in (C) treated for 24 hrs with mizoribine (2 μM), alone and in combination with the ATR inhibitor AZD6738 (2 μM) added for the last 6 or 12 hrs as indicated. Cells were pulse labeled with EdU (10 μM) for the last two hours, followed by DAPI staining. **(F)** Representative images. Scale bar = 50 μm. **(G)** EdU-positive cells were quantified and graphed as percentage of the total cell population. The 12 hr samples were collected in the same experiment as Figure 5C,D, thus data shown for the vehicle, mizoribine alone, and 6 hr AZD6738 samples are the same as in Figure 5C,D, shown again here for comparison to the 12 hr samples.

Data are presented as mean of three biological replicates ± SEM. *$p<0.05$ by Welch's t-test.

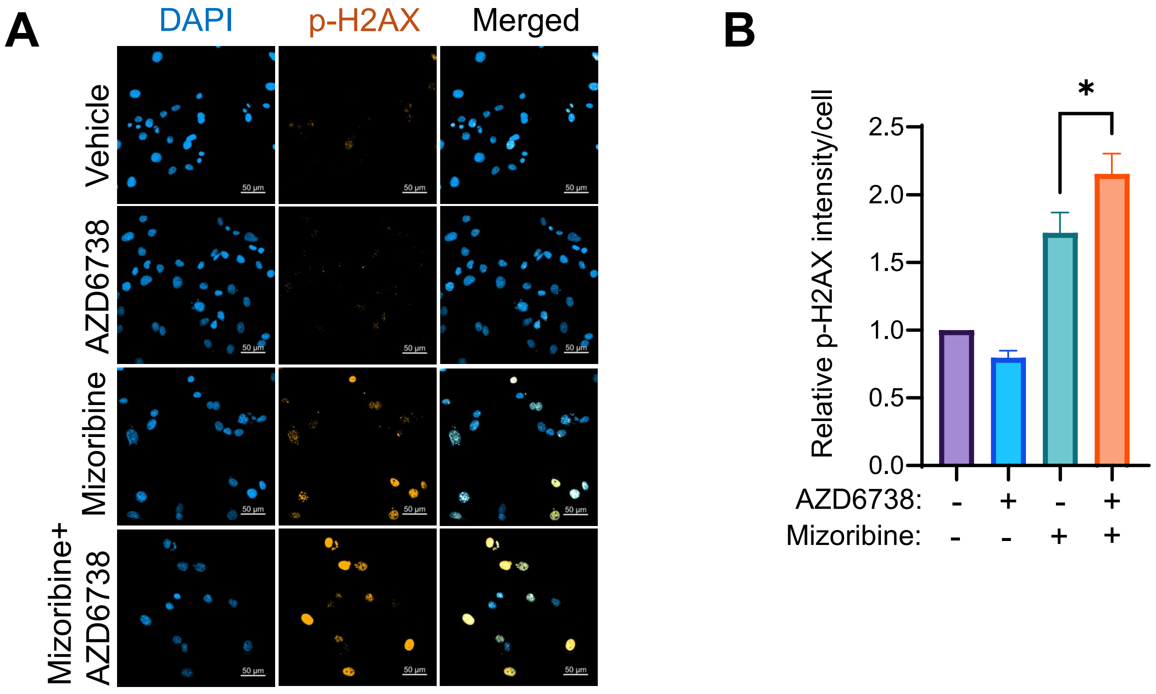

A
DAPI
p-H2AX
Merged
Vehicle
AZD6738
Mizoribine
Mizoribine+
AZD6738
50 µm
B
Relative p-H2AX intensity/cell
2.5
2.0
1.5
1.0
0.5
0.0
*
AZD6738: - + - +
Mizoribine: - - + +

**Supplemental Figure 5. Supplemental data supporting Figure 6**

**(A, B)** *Tsc2*$^{-/-}$ *Trp53*$^{-/-}$ MEFs treated with mizoribine (3 μM) for 48 hrs, alone and in combination with the ATR inhibitor AZD6738 (2 μM) added for the last 12 hrs, followed by immunofluorescence staining for p-H2AX S139 (p-H2AX). **(A)** Representative images, scale bar = 50 μm. **(B)** p-H2AX staining intensity per cell was quantified and graphed as mean ± SEM relative to control. n = 275 cells per condition. *p<0.05 by Welch's t-test.
